# Human conflict with wild carnivores and herbivores: a comparison of socio-economics, damage and prey preference of wild carnivores

**DOI:** 10.1101/2022.08.22.504800

**Authors:** Abhijeet Bayani, Nikhil Dandekar

## Abstract

Human-wildlife conflict (HWC) at the fringes of protected areas is a major concern in the conservation biology. Although damage caused by carnivores and herbivores may vary in their magnitude, more attention has been given to carnivores due to various reasons. In Tadoba-Andhari Tiger Reserve (TATR) we compared economic dependence of locals on agriculture and livestock independently and found that income from livestock is only additional, whereas agriculture is the major source of livelihood. We also studied the relative abundance of wild herbivores and diet of tiger, leopard and sloth bear and found that these carnivores have largely been dependent on chital, nilgai and wild pig despite livestock population being higher. We found HWC mitigation in TATR effective but biased towards big cats while the damage in main livelihood (agriculture) being higher. We recommend higher attention to crop-raiding problem for the successful wildlife conservation in central India.

## Introduction

Co-existence of humans and the large wild mammalian species does not always remain harmonious especially at the fringes of protected areas leading to human-wildlife conflict (HWC). Human casualties due to carnivores (and by some mega-herbivores like elephant) is the most severe case of HWC. All over the globe, it has rightfully gained enough attention and it has been studied extensively and carefully. On the other hand, HWC may turn equally severe when residents near protected areas face substantial loss to their livelihood sources (Treves et al., 2006; 2020; Zimmermann et al., 2020) especially to the livelihood from livestock and from the agriculture. In many cases, the damage caused by carnivores (to livestock) and by the herbivores (to agriculture) is of substantial magnitude and it causes grave resentment among the victims (Marchini, 2014; Zimmermann et al., 2020) leading to retaliatory killing of the wildlife. Such a reaction to livelihood loss has been identified as one of the most detrimental causes for the loss in wildlife. To avoid such a resentment, there are several legal and government-approved policies and methods developed to compensate for such livelihood loss faced in either of the cases. The compensation for loss caused by wildlife (monetary or otherwise) has been observed to be efficient in reducing the antipathy towards the wildlife and has been effective in reducing the retaliatory killing as well (e.g. Groom & Harris, 2008; Johnson et al., 2018).

Nevertheless, there are many subtle differences in the compensation methods, approach, attitude, damage estimation methods, and the amount paid for the loss when compared between the damage caused by carnivores and the damage caused by the herbivores (Woodroffe et al., 2005; Douglas & Verissimo, 2013, Nyhus et al., 2003; Bulte & Rondeau, 2005; Bayani et al., 2016; Dickman, 2010; Marchini, 2014; Verschuren et al., 2020). There are two main components to the system of compensation, namely, the victims and the organization that pays the actual compensation. Thus, the differences in the overall methodology involved in compensation should be viewed from two perspectives (1) Victims’ perspective and (2) Policy makers’ perspective.

1. Victims’ perspective: A victim pursues the compensation when the procedure is easy, and the benefit is higher than the loss. Victim may also pursue the compensation when the livelihood dependence on the source (livestock and agriculture) is excessive and the loss is enormous. In the later case, a victim would pursue the compensation despite the complexity of the procedure. It is however, imperative that if the procedure is long, tedious and not well assured (for receipt of compensation amount), the victim may not pursue it. A victim may also not pursue it if the calculation and measurement of loss is ambiguous and not transparent enough to be trusted, and/or the compensation amount to be obtained is not substantially higher than what the victim may have to spend for the procedure.
2. Policy makers’ perspective: A loss caused by wildlife is expected to be compensated without being biased by the amount to be paid, amount of loss, or the economic dependence (on the source to which the loss has occurred) of the victim. If the possibility of resentment against wildlife is higher, policy makers would not risk it and thus the compensation claim would be passed more actively, the amount of compensation would be elevated, or the procedure would be made easier. Thus, it may happen that certain claims or types of claims would have higher priority than the others. For instance, damage caused by tiger maybe prioritized over the damage caused by wolf or damage by elephants maybe taken more seriously than that caused by wild pigs. Such compensation assumes the quick and unbiased measurement of loss. Nevertheless, if the damage estimation itself is tedious or ambiguous, policy makers remain equally helpless.

In case of damage caused by carnivores, compensating for the number of livestock individuals killed or injured is the unbiased method which is consented by compensating organization and victims both. In case of agricultural loss, when an entire standing crop is totally trampled becomes easy to measure and compensate for (assuming that the expected crop yield is fixed and agreed by both parties). In all these various scenarios of damage and compensation. One important component is either neutralized or not considered, which is the actual economic dependence. Which may make the compensation system more lucrative.

During our initial studies (Bayani et al. 2016) in Tadoba-Andhari Tiger Reserve (central India) we found that damage estimates of crop loss and the compensation for the same were neither well-formulated nor pursued by claimants (the farmers) nor by the authorities. On contrary, the loss in the livestock was well documented, claimed more actively and was compensated without much technical difficulty. There could be several factors responsible for such a difference. Considering the two perspectives described above we put forth 4 hypotheses to test (1) dependence of large carnivores on livestock is higher (thus carnivores kill livestock more frequently) and to avoid any resentment against wildlife such loss is actively compensated by government; (2) the economic loss due to the livestock damage is higher and hence it is compensated actively; (3) economic dependence of locals on livestock is higher and hence it is pursued by claimants more rigorously, (4) as compensation to livestock damage is easier and quicker hence it is pursued by claimants more actively irrespective of the livelihood dependence on it.

To test these hypotheses, we collected data on (i) income from agriculture and livestock, (ii) frequency and amount of damage to agriculture and livestock due to wildlife, (iii) cases of retaliatory killing, (iv) cases of compensation for livestock and agriculture. The data on income were collected through well-designed socio-economic surveys; the data on damage in agriculture and livestock were obtained from detailed sampling in the field, case studies and partly from the questionnaires; data on retaliatory killing and compensation were obtained from case studies. Additionally, we studied the dependence of carnivores on different available prey species through sampling in the field and the scat analysis. The details of damage and its estimation are already published and can be obtained from Bayani et al. 2016 and Watve et al. 2016.

## Methods

Our study commenced between year 2012 and 2015 at the western boundary buffer of Tadoba-Andhari Tiger Reserve (TATR) situated in the eastern part of Maharashtra state in central India. We divided the sampling and data collection for this study into four categories: (1) questionnaire or socio-economic surveys, (2) livestock predation case studies, (3) herbivore population estimation, and (4) scat collection and analysis. Sampling for all these was done simultaneously throughout the study period. The case studies were only opportunistic based on availability to the process of compensation claims.

### Questionnaire surveys

We selected 12 villages in the western boundary buffer to conduct socio-economic surveys. Before beginning the surveys, we always met the village head and obtained oral consent to conduct interviews. The respondents were chosen at random but we cautiously avoided interviewing more than one person from one family to prevent redundancy in the data. The interviews were conducted in local language with the help of local farmers who were trained by us before beginning the surveys. The questionnaire forms were printed in local language and script (*Marathi*) so that respondents could read and understand the questions and the reported responses (written by authors). We asked factual questions about their sources of income, basic agricultural and livestock holding practices, grazing practices, and attitude towards wildlife.

### Herbivore population study

In the western boundary buffer, we selected a 40 sq. km area to conduct wild animal population surveys, which was devoid of villages or farms. We followed distance sampling using standard line transect method (Buckland et al., 1993) to estimate density and relative abundance of wild herbivore and livestock population in the study area. The sampling was done in all three seasons i.e. Monson (July-October), winter (November-February), and summer (March-June). Each transect line was 2 km in length laid randomly in the study area and were walked in the daytime. We used handheld GPS to keep track of distance and direction and used rangefinder to measure the sighting distances. We walked 227 transects in total covering total transect length 454 km. The study area was heavily used for grazing by livestock. As our pilot studies had indicated that carnivores also prey on the livestock, we collected data on the population of livestock during this sampling counted in the same manner as the wild herbivores on the transects.

### Scat collection and Analysis

For carnivore scat collection we walked various pre-identified routes and pathways in the entire study area. We collected only tiger, leopard and sloth bear scats (wild dog is another major predator in TATR but it did not frequent enough in the buffer areas and hence not enough scats could be collected during the study period). The big cat scats that we collected were mostly that of tiger, although we did not attempt to distinguish tiger scat from that of leopard. However, based on our long-term observations in the area, knowledge of local people and forest guards, it was evident that leopard did not frequent that area substantially and hence its representation in the analysis is expected to be negligible.

The scats were collected in plastic Ziploc bags and later preserved in 70% ethanol. We followed Bahuguna et al. (2012) for scat processing, medullar slide preparation and identification of prey species using the hair obtained from the collected scat samples.

## Results

### Agriculture and livestock holding

We interviewed 439 local residents distributed across 12 villages independently. All the respondents claimed to have three major income sources *viz*. agriculture, livestock holding and labor work on daily wages. All the respondents were found to utilize these three income sources except for about 3% (13 out of 439) of them that claimed being entirely dependent on the seasonal labor work. Farmers in TATR practice agriculture in the two crop seasons i.e. monsoon (locally known as *kharif* i.e. June to October) and winter (locally known as *rabi* i.e. November to February). They may switch to labor work or any other non-agricultural activity in non-cropping season i.e. summer (March to June). The income per year from agriculture alone was found to be INR 40337±49321 (Mean±SD) per family (Range: INR 3000 to INR 600000), from livestock INR 16767±15959 (Mean±SD) per family (Range: INR 500 to 90000), whereas the income from labor work was not fixed nor was evaluated. It was treated only as a surplus income and respondents did not keep any track on this. However, it did not exceed INR 10000 per year per family. It is important to note that about 65% respondents did not depend economically on livestock at all although they held livestock with them.

Livestock holding included cattle (cows and bullocks), buffalo, goat and sheep. 97.95% farmers had at least one of the above listed livestock. 4.32% people owned buffalos and average adult individuals per family was 2.73±1.6 (range 1 to 6); 52.6% families have these buffalos passed to them traditionally (from earlier parental generation) which they had not purchased, whereas 47.3% families purchased at least one adult buffalo to utilize it for the milk production as an additional income source. Purchase cost of buffalo ranged between INR 2500 and INR 30000, that provided an annual income per family ranging between INR 1000 and INR 24000. Per annum cost of maintenance of buffalo ranged between INR 100 to INR 500 per adult individual. This included vaccination, fodder, and other.

Bullocks are essential for agricultural activities and hence all the farmer families (except the ones who depend on labor work entirely), owned at least one adult bullock. Number of adult bullocks per family ranged between 1 and 8 (on an average 2.7±1.26 individuals per family), 52.68% people had purchased at least one bullock whereas 47.7% families had obtained them traditionally as a family asset. Purchase price per pair of bullocks ranged between INR 750 to INR 100000 (19006.2±16201.8). Bullocks did not provide any additional income to farmers although some farmers, especially the ones those owned only one bullock individual borrowed/rented another bullock from another farmer on a per day rent i.e. INR 100 to INR 300 per day. Maintenance cost per individual was found to be ranging between INR 100 to INR 2000 per annum.

Number of adult cows per family ranged between 1 to 15 (average 3±2). About 64% families obtained cows as family asset and had not purchased them separately. Rest proportion (36%) had purchased it. The purchase cost per individual ranged between INR 300 to INR 6000. The cows provided an additional income to farmers that ranged from INR 500 to INR 50000 per annum. Maintenance cost per individual per annum was INR 100 to INR 1500.

33.7% farmers owned goat and sheep, although number of sheep was negligible (hence not counted here). Average number of adult goats per family was 3.57±5.38 (1 to 52). As much as 43.9% farmers did not purchase the goat but got them as family asset, whereas 56% farmers have purchased at least one pair of goats. Purchase cost of goat per individual ranged from 300 to 4000, whereas maintenance cost per individual did not exceed 50 per individual per annum. Goat was not considered a preferred secondary source of income but was rather raised for own consumption. Adult goat was occasionally sold out to other thus income from goat was not consistent for farmers.

### Grazing practices

Buffalos and bullocks were almost always taken out for grazing on farms or remained close to village, which were herded and taken care otherwise by the owners themselves irrespective of the season. Cows, young bullocks, goat and sheep of a village were handed over for grazing to a common shepherd belonging to a same village. All these three livestock species were herded separately and were never found to be mixed. The grazing grounds and grazing practices changed as per the season. In Summer the cow herds were set rather scattered and free ranging without any active supervision (85.38%) or fed/grazed at home (15.08%). Cow herds in summer formed smaller scattered herds and were distributed at grazing areas rather more randomly than any other season. Goat and sheep were fed at the owners’ residence and were seldom set free out to grazing areas in summer.

In monsoon and winter, all livestock species (except bullocks and buffalo) were taken to protected areas and abandoned farms. The herds were herded by one or two shepherds and who always made tight herds of large number of individuals aggregated at one area. This is also evident in population estimation data. Through the interviews of farmers, herders and observations at the grazing grounds it was understood that livestock remained at the grazing grounds in protected areas and for at least 5 hours between 930h to 1630h.

The herders of all villages and the livestock owners were aware of grazing grounds and presence of at least one of the three carnivores *viz*. tiger, leopard and sloth bear in the areas where the herds were taken for grazing. Some herders even had claimed to have seen wolf and wild dogs which were potential threats to livestock.

All the villages are situated within 400 m to 2 km distance from the tiger reserve boundary. Thus, all the farmers had constructed greater livestock protection means at their residence. Close to 88% farmers had at least one cattleshade made up of bamboo and timber harvested from forest. Rest others had bamboo and wired fence as an additional protection measure.

### Livestock depredation and compensation

There were total 103 instances of livestock depredation during the study period. Out of these, partial information on 51 cases could be obtained from the questionnaire interviews whereas other 52 cases could be studied on ground completely (from tracing the killed livestock to the receipt of compensation amount to individual owners). When we interviewed locals, it turned out that as much as 23.4% families had lost at least one livestock species (cow, sheep or goat) to carnivores in previous three years. 73.78% (76 out of 103) cases were claimed for the loss and received with the compensation. None of the respondents claimed to have lost more than one livestock individual during the study period. From the 52 completely studied cases of livestock predation, we found none being poisoned or tampered in any way indicating any retaliatory killing. Informal interviews with the forest guards also indicated that there were no cases of retaliatory killings during the study period or in the previous 5 years.

As per the government approved procedure for the compensation claim, the carcass of killed livestock is needed for the examination. If in case the killed livestock remains untraceable or unobtainable for examination, claim could not be made. During our study, we found that most of goat and sheep carcasses could not be traced and hence could not be claimed for the compensation. In case when they were obtained back, the compensation amount was found to be ranging between INR 200 and 1000 per claim (average purchase cost being INR ∼1700 per adult goat). In case of cattle (calves, young bullocks, and young and adult cows) most of the carcasses could be retrieved and claimed for the compensation. The compensation amount ranged between INR 4000 and 8000 (while the average purchase cost was INR ∼3700). There were no instances adult bullocks found to be depredated by any wild carnivore during the study period.

### Diet of large carnivores (tiger, leopard and sloth bear)

The carnivores (tiger and/or leopard) predated on total 14 herbivore species (Figure 1A, B, C). It was found that presence of livestock in their diet was less than 15% in any season. This means that tiger and leopard were largely dependent on wild herbivores for their foraging (Figure-1A, B, C). Wild dogs (dholes) were seldom seen in the peripheral area and there were no reported instances of them feeding on livestock. Wolf was seen only once in the study area that predated on 2 goats in one day. We did not include this data in our analysis. Sloth bear is another frequent carnivore in the study area. It is reported that Sloth Bear may scavenge on the carcasses or leftovers from tiger/leopard kills (Joshi et al., 1997). We analyzed its scats and found no traces of any herbivore (wild or livestock) in its diet.

**Figure 1.**
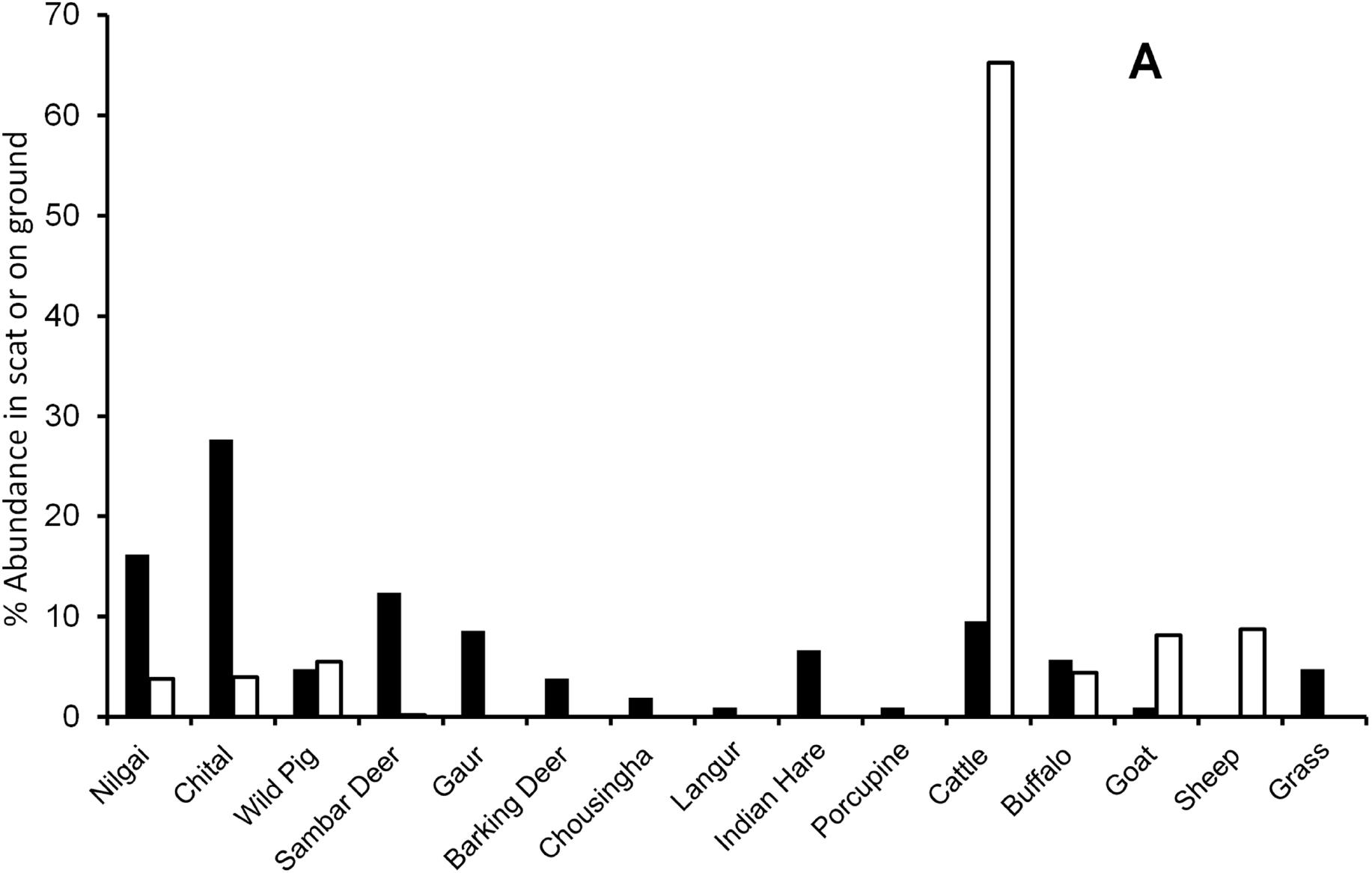
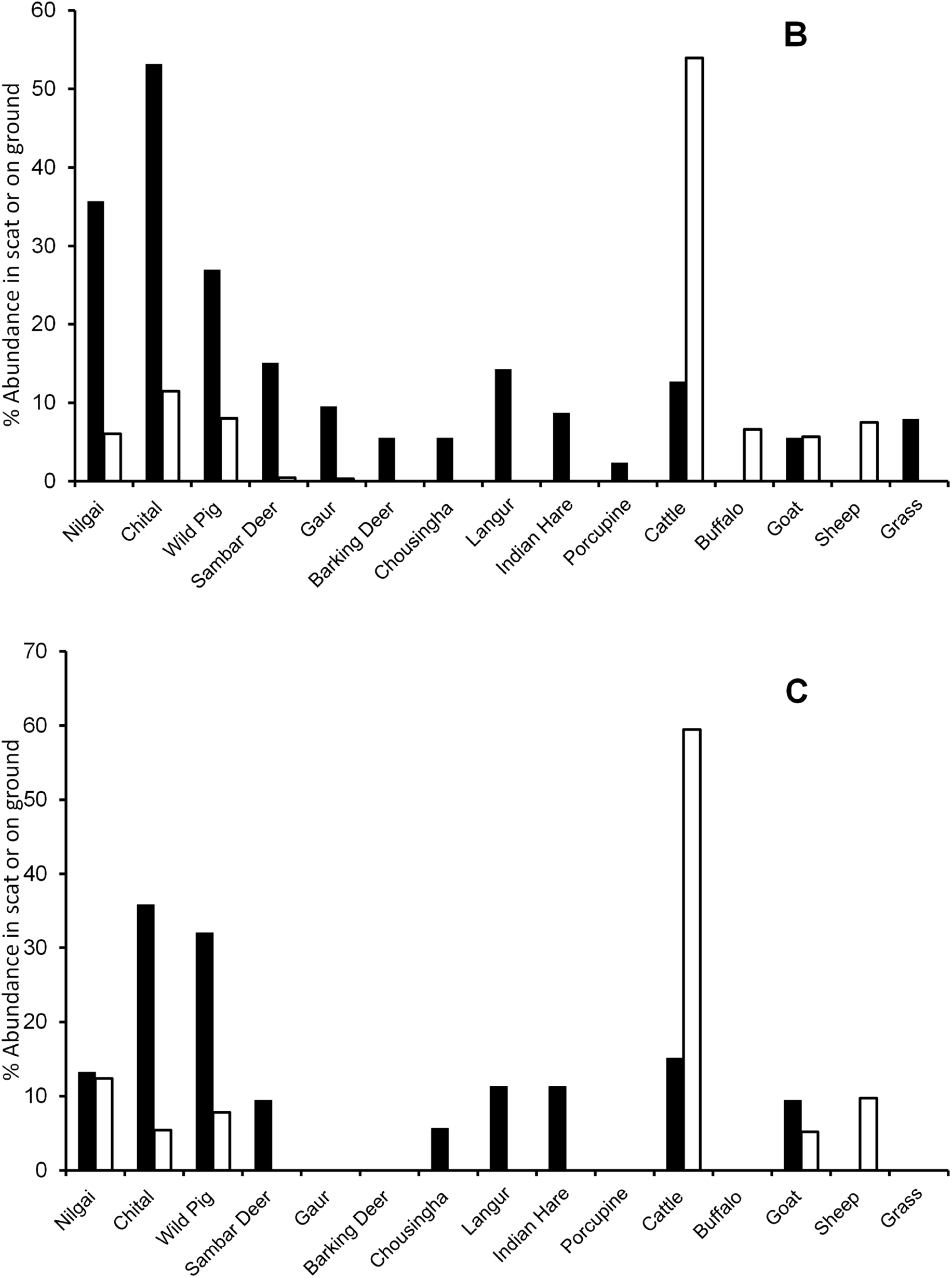
Comparison of wild herbivores and livestock abundance on ground (%, per sq. km, clear empty bars) with the % abundance of different herbivore species observed in the scat analysis (solid black bars). Abundance of Langur, Indian Hare, Porcupine and grass was not evaluated during the transect surveys but were evaluated from the scat analysis, thus absence of empty bars for those species does not indicate their absence from the field or on transects. A, B, and C are three different graphs for three seasons: (A) monsoon, (B) winter, (C) summer.

### Herbivore population in the peripheral areas of study area

We found that in all the seasons, population of livestock was much higher that the wild herbivores (Figure-1A, B, C). Among livestock, cattle population was seen to be highest; whereas Nilgai (*Boselaphus tragocamelus*), Chital (*Axis axis*) and Wild Pig (*Sus scrofa*) were the most abundant wild herbivore species. It should be noted that the livestock population was much scattered in the summer than in the other two seasons owing to the herding strategies explained above. It is also interesting to see that despite high population of cattle, chital was the main food item found in the scat analysis. This clearly indicates that big cats in peripheral areas of TATR prefer chital over all other available herbivores irrespective their relative abundance.

### Comparison of loss in livestock and in agriculture during the study period

We found that during the study period the absolute average loss in the livestock per capita is about 5000 INR (only considering the locals who have non-zero income from livestock), whereas average loss in agriculture is about 20000 INR. We observed an exactly opposite trend in the gain from compensation. The gain from compensation for livestock ranged between 4000 INR and 8000 INR, and the same for agriculture ranged between 3000 INR and 5000 INR. The deficit in the loss and compensation is orders of magnitude different, making it more severe for the agriculture (Figure-2). It can also be seen from Figure-2 that ∼73% loss is compensated whereas the same for agriculture is only 21%. It is evident from the analysis that locals economically depend more agriculture than livestock but despite having more loss in the agriculture, compensation is not actively pursued. We found that the methods of crop loss compensation were more tedious and less rewarding than those for the livestock loss (Watve et al. 2016 discusses the reasons for the same more rigorously).

**Figure 2.**
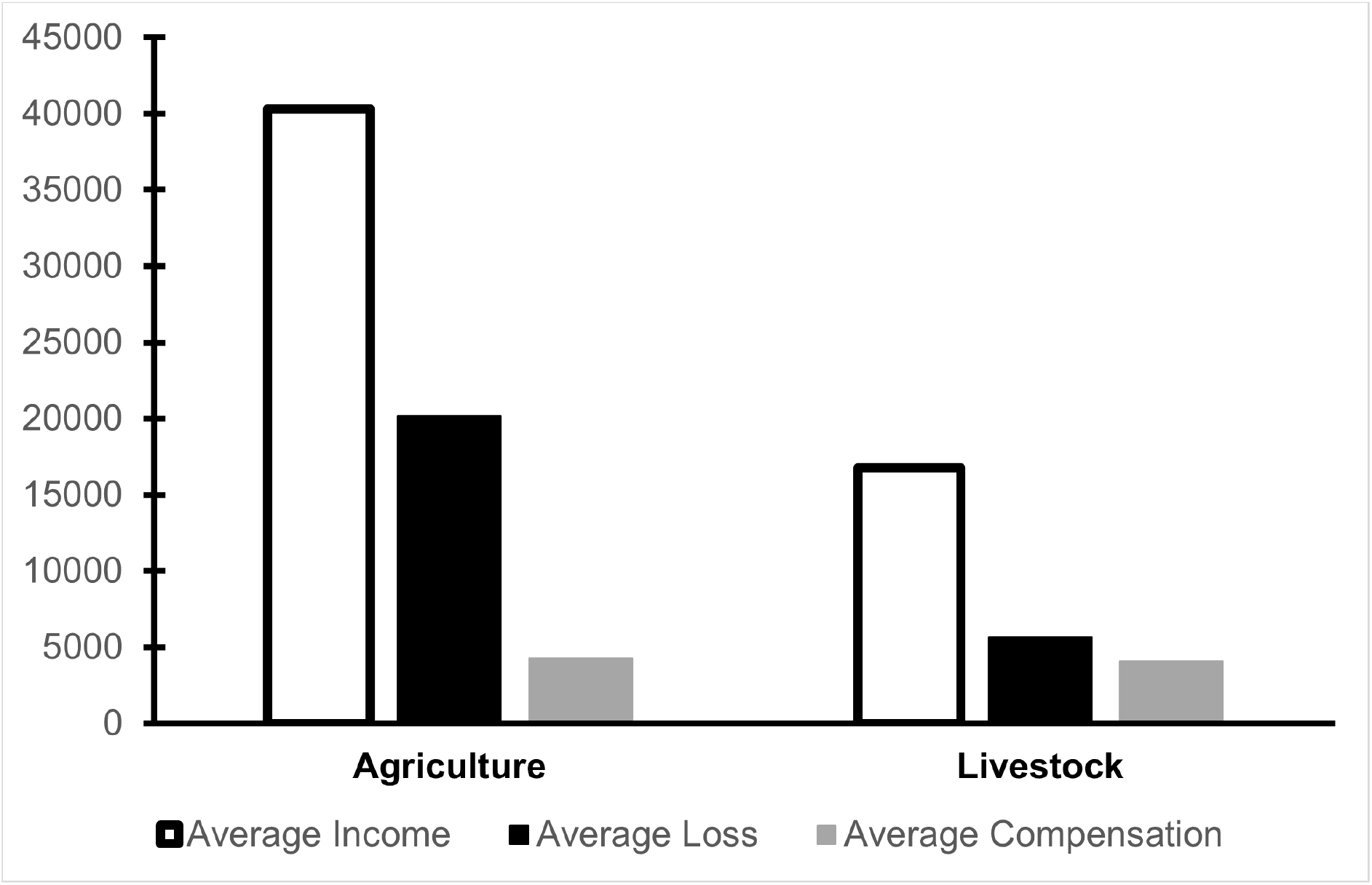
Comparison of average income, average loss and average compensation per capita in agriculture and livestock in TATR. The numbers on Y-axis are in INR. Empty/clear bars represent Average income (INR), solid black bars indicate average loss (INR), grey bars indicate average compensation received (INR).

We fail to reject the hypothesis that ‘compensation to livestock damage is easier and quicker hence it is pursued by claimants more actively irrespective of the livelihood dependence on it.’ We, however, reject the following two hypotheses, namely: ‘the economic loss due to the livestock damage is higher and hence it is compensated actively’; and ‘the economic dependence of locals on livestock is higher and hence it is pursued by claimants more rigorously’. Our studies also clearly indicate that the dependence of large carnivores on livestock is lower, nevertheless, it is highly likely that loss in livestock is compensated well to avoid any resentment against wildlife.

## Discussion

It is generally assumed and also observed that livestock is an easy and abundant prey and hence wild carnivores are more likely to hunt the livestock (e.g. Linnell et al., 1999; Kolipaka et al., 2017; Soofi et al., 2019). Our data differs from this perception showing despite high abundance of livestock in the areas where carnivores are frequent, they still showed preference to wild herbivores. Local community perhaps know this fact through their experience or the traditional knowledge. It is also highly likely that the perceived risk of livestock being predated on grazing ground among the local people is low because (i) livestock is taken for grazing to the areas that the carnivores frequented, (ii) the livestock depredation is temporally stochastic, and (iii) the compensation is readily available with fairly easy and community-friendly process (also see Watve et al., 2016). It should also be noted that livestock keeping although is necessary for the farming operations, locals are not dependent on it economically. It is therefore plausible that the income from livestock including that obtained through compensation could only be surplus. This also indicates that the conflict mitigation (at least the compensation) in our study area must have been effective having no retaliatory killings of any wildlife observed.

It is quite evident that agriculture is the main occupation of locals. Damage from the wild herbivores to the crops causes 50-70% economic loss per year yet the crop damage compensation methods are more tedious resulting into no gain for the farmer communities (see Bayani et al., 2016). There are only a few attempts to address this problem (e.g. Watve et al. 2016, Joshi et al., 2021) throughout the central India. Although severity of the conflict with both wild carnivores and herbivores still needs better evaluation, it can be seen that magnitude of loss in agriculture is orders of magnitude higher than that in livestock. We think that to achieve better and active involvement of locals in the wildlife conservation, livelihood of locals should be secured first.

TATR is the major conservation area in India. There has been substantial sensitization among local people about the conservation and good involvement of them in the conservation activities. At TATR, the agriculture being the major livelihood source, it should be secured first to obtain a continued support from the locals. We suggest that there needs more research and better management or mitigation strategy to reduce the crop depredation in central India.

## Acknowledgements

We thank Department of Forest, Maharashtra, India and Tadoba-Andhari Tiger Reserve for the support and work permits. We greatly acknowledge Indian Institute of Science Education and Research Pune and Abasaheb Garware College, Pune for providing us the facility to work. We thank Prof. Milind Watve (Scientist, Deenanath Mangeshkar Hospital and Research Center, Pune) for their guidance during the research and letting us use the field station at TATR.

